# Ebola Virus Sequesters IRF3 in Viral Inclusion Bodies to Evade Host Antiviral Immunity

**DOI:** 10.1101/2023.04.20.537734

**Authors:** Lin Zhu, Jing Jin, Tingting Wang, Yong Hu, Hainan Liu, Ting Gao, Qincai Dong, Yanwen Jin, Ping Li, Zijing Liu, Yi Huang, Xuan Liu, Cheng Cao

## Abstract

Viral inclusion bodies (IBs) commonly form during the replication of Ebola virus (EBOV) in infected cells, but their role in viral immune evasion has rarely been explored. Here, we found that interferon regulatory factor 3 (IRF3), but not TANK-binding kinase 1 (TBK-1) or IκB kinase epsilon (IKKε), was recruited and sequestered in viral IBs when the cells were infected by EBOV transcription- and replication-competent virus-like particles (trVLPs). NP/VP35-induced IBs formation was critical for IRF3 recruitment and sequestration, probably through interaction with STING. Consequently, the association of TBK1 and IRF3, which plays a vital role in type I interferon (IFN-I) induction, was blocked by EBOV trVLPs infection. Additionally, IRF3 phosphorylation and nuclear translocation induced by Sendai virus (SeV) or poly(I:C) stimulation were suppressed by EBOV trVLPs. Furthermore, downregulation of STING significantly attenuated VP35-induced IRF3 accumulation in IBs. Coexpression of the viral proteins by which IBs-like structures formed was much more potent in antagonizing IFN-I than expression of the IFN-I antagonist VP35 alone. These results suggested a novel immune evasion mechanism by which EBOV evades host innate immunity.

**Impact statement:** Ebola virus VP35 protein evades host antiviral immunity by interacting with STING to sequester IRF3 into inclusion bodies and inhibit type-I interferon production.

## Introduction

Ebola virus disease is the deadliest infectious disease caused by infection with Ebola virus (EBOV), an enveloped, nonsegmented negative-sense RNA virus (Feldmann, Jones, Klenk, & Schnittler, 2003). The 19-kb viral genome comprises seven genes encoding the nucleoprotein (NP), virion protein 35 (VP35), VP40, glycoprotein (GP), VP30, VP24 and RNA-dependent RNA polymerase (L) (Mahanty & Bray, 2004). Inclusion bodies (IBs) that form in EBOV-infected cells are specialized intracellular compartments that serve as sites for EBOV replication and the generation of progeny viral RNPs (Hoenen et al., 2012; Nanbo, Watanabe, Halfmann, & Kawaoka, 2013). In IBs, the EBOV genome is replicated and transcribed by viral polymerase complexes (Misasi & Sullivan, 2014). VP35 serves as a cofactor of RNA-dependent RNA polymerase and contributes to viral replication by homo-oligomerization through a coiled-coil domain (Reid, Cardenas, & Basler, 2005) as well as through its phosphorylation and ubiquitination, which was recently discovered (van Tol et al., 2022; Zhu et al., 2020).

Innate interferon responses constitute the first lines of host defense against viral infection. Retinoic acid-inducible gene I (RIG-I)-like receptors (RLRs), including RIG-I and melanoma differentiation-associated protein 5, play pivotal roles in the response to RNA virus infection. After the recognition of RNA virus infection, RIG-I is recruited to the mitochondrial antiviral adaptor protein (MAVS) through the caspase activation and recruitment domain. The activation of MAVS recruits multiple downstream signaling components to mitochondria, leading to the activation of inhibitor of κ-B kinase ε (IKKε) and TANK-binding kinase 1 (TBK1), which in turn phosphorylate IFN regulatory factor 3 (IRF3). Phosphorylated IRF3 forms a dimer that translocates to the nucleus, where it activates the transcription of type I interferon (IFN-I) genes (Fitzgerald et al., 2003; Liu et al., 2015).

To promote viral replication and persistence, viruses have evolved various strategies to evade or subvert host antiviral responses. For example, severe fever with thrombocytopenia syndrome virus (SFTSV) has developed a mechanism to evade host immune responses through the interaction between nonstructural proteins and IFN-I induction proteins, including TBK1, IRF3 and IRF7 (Hong et al., 2019; Lee & Shin, 2021; Ning et al., 2014; Wu et al., 2014), sequestering them inside SFTSV-induced cytoplasmic structures known as IBs. In addition to inhibiting IFN-I induction, SFTSV nonstructural proteins can hijack STAT1 and STAT2 in IBs to suppress IFN-I signaling (Ning et al., 2015). These studies highlight the role of viral IBs as virus-built “jails” that sequester some crucial host factors and interfere with the corresponding cellular processes.

EBOV uses various approaches to evade the host immune response, including antagonizing IFN production, inhibiting IFN signaling, and enhancing IFN resistance (Basler et al., 2000; McCarthy et al., 2016; Reid et al., 2006). VP35 is an IFN-I inhibitor that antagonizes host innate immunity by interacting with TBK1 and IKKε (Basler et al., 2003; Prins, Cardenas, & Basler, 2009), suppressing RNA silencing and inhibiting dendritic cell maturation (Haasnoot et al., 2007; Yen, Mulder, Martinez, & Basler, 2014). Here, we report that viral IBs in EBOV transcription- and replication-competent virus-like particles (trVLPs)-infected cells appear to play a role in immune evasion by sequestering IRF3 into IBs and preventing the interaction of IRF3 with TBK1 and IKKε.

## Results

### IRF3 is hijacked into cytoplasmic IBs in EBOV transcription and replication-competent virus-like particles infected cells

When HepG2 cells were infected with EBOV trVLPs (Hoenen, Watt, Mora, & Feldmann, 2014), which authentically model the complete virus life cycle, IBs with a unique structure and viral particles formed in the cytoplasm (Fig. 1 – figure supplement 1A-B). Surprisingly, we found that a substantial percentage of endogenous IRF3 was trapped in viral IBs in EBOV trVLPs-infected cells with large IBs (Fig. 1A and 1B), while no detectable TBK1 or IKKε, the essential upstream components of IRF3 signaling (Fitzgerald et al., 2003), was sequestered in the viral IBs (Fig. 1C-F). These results suggested that IRF3 was specifically compartmentalized in viral IBs, and this compartmentalization spatially isolated IRF3 from its upstream activators TBK1 and IKKε.

**Fig. 1.**
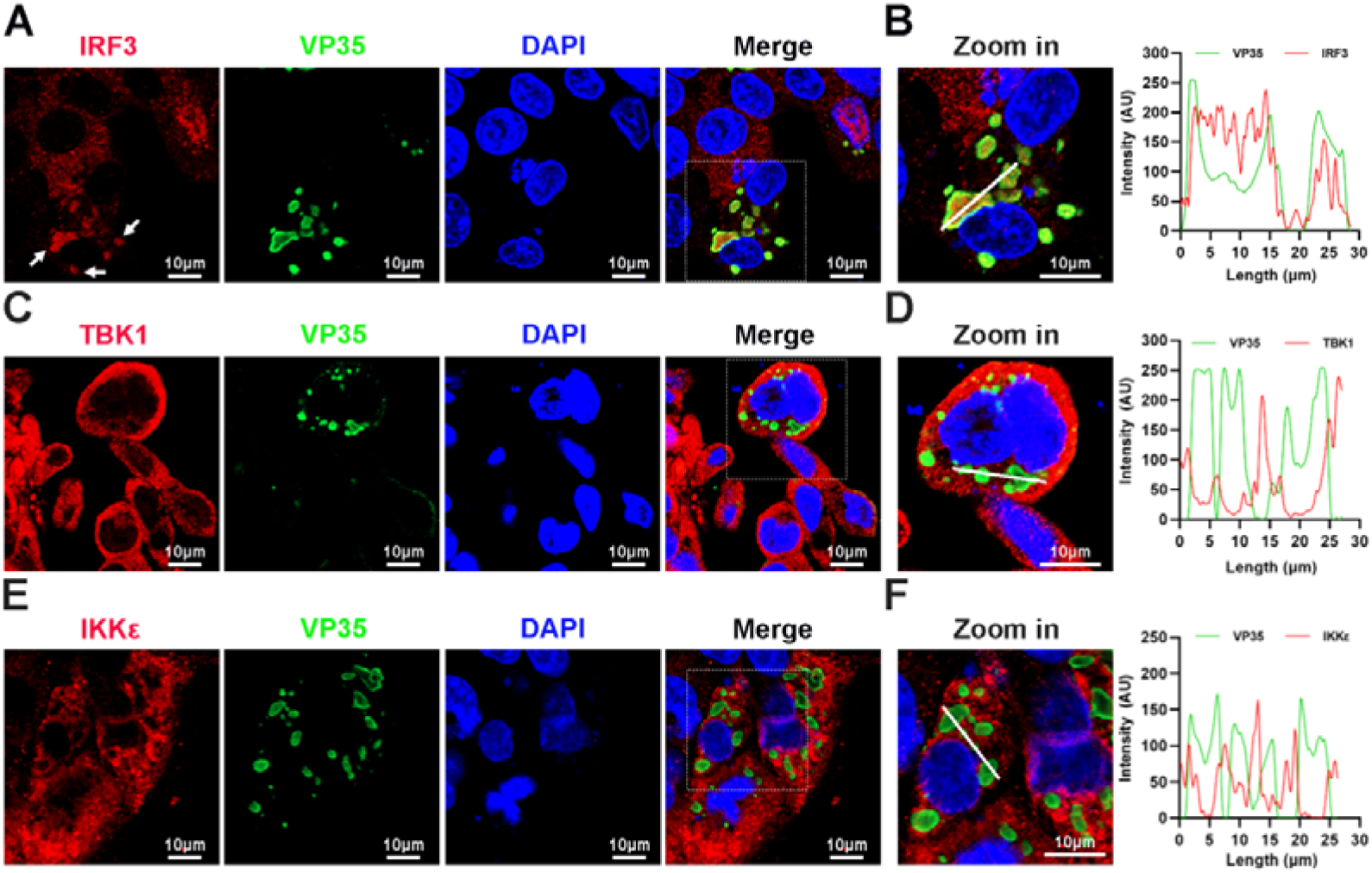
IRF3, but not TBK1 and IKKε, is sequestered into viral inclusion bodies (IBs) upon EBOV trVLPs infection. (A), HepG2 cells infected with the EBOV trVLPs were immunostained with anti-IRF3 (red) and anti-VP35 (green) antibodies. Nuclei were stained with DAPI (blue), and images were obtained using a Zeiss LSM 800 Meta confocal microscope. White arrows: IRF3 in IBs. (B), The left panel shows a magnified image of the IBs boxed in the merged panel of (A). The graphs (right panel) show the fluorescent intensity profiles along the indicated white lines drawn across one or more IBs. (C and E), HepG2 cells infected with the EBOV trVLPs were immunostained with anti-TBK1 (red in (C)) or anti-IKKε (red in (E)) and anti-VP35 (green in (C and E)) antibodies. Nuclei were stained with DAPI (blue), and images were obtained using a Zeiss LSM 800 Meta confocal microscope. Scale bar, 10 μm. (D and F), The left panel shows a magnified image of the IBs boxed in the merged panel shown in (C) and (E). The graphs (right panel) show the fluorescent intensity profiles along the indicated white lines drawn across one or more IBs.

The sequestration of IRF3 in IBs was further investigated at different hours post infection (hpi) of EBOV trVLPs. Detectable IRF3 puncta colocalized with viral proteins were apparent at 36 hpi in infected cells and correlated significantly with the size and shape of the viral IBs (Fig. 2A and 2B). As the size of IBs increased at 48 hpi, nearly all IRF3 colocalized with viral IBs, whereas the IRF3 distribution was completely different in the uninfected cells nearby (Fig. 2A and 2B). Using a fluorophore line of interest analysis, we assessed the intensity profiles of cytoplasmic IRF3 intensity in IBs as well as the increase in the diameter of the aggregates (Fig. 2C). As infection proceeded, the intensity of the IRF3 signal in the puncta increased as the level of cytoplasmic-dispersed IRF3 decreased (Fig. 2A), indicating that the total amount of IRF3 in the cells did not dramatically change during infection (Fig. 2D and 2E) and that only its subcellular localization changed. Taken together, the results above showed that IRF3, but not TBK1 or IKKε, was sequestered in viral IBs.

**Fig. 2.**
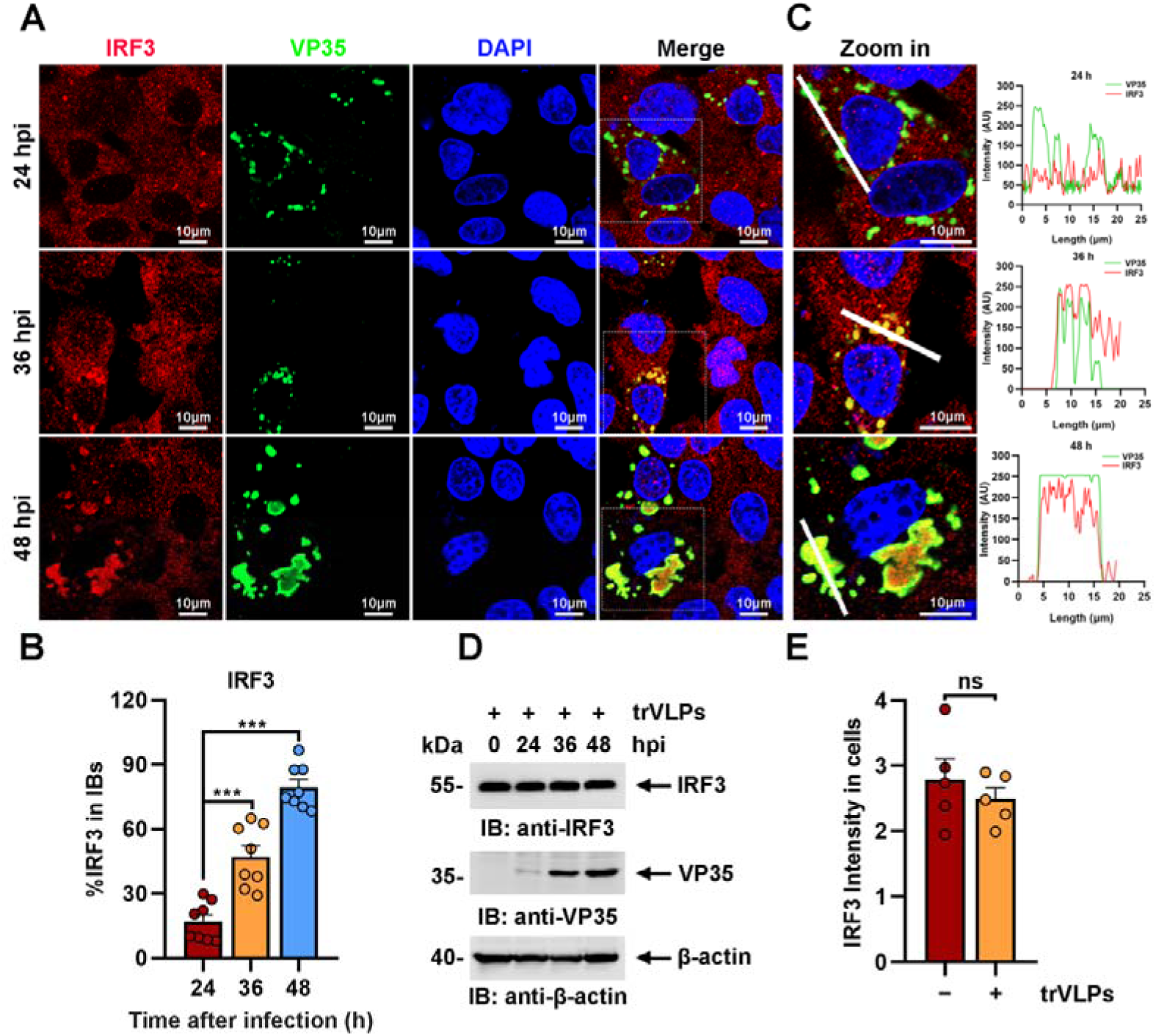
EBOV trVLPs induce the recruitment of IRF3 into intracytoplasmic IBs. (A), HepG2 cells were infected with EBOV trVLPs. At the indicated time points after infection, cells were fixed and immunostained with anti-IRF3 (red) and anti-VP35 (green) antibodies. Nuclei were stained with DAPI (blue), and images were obtained using a Zeiss LSM 800 Meta confocal microscope. Scale bar, 10 μm. The data from two independent replicates are presented. (B), The percentage of IRF3 distribution in IBs at different time points in cells infected with EBOV trVLPs (A) was analyzed using the R programming language. The intensity of IRF3 in 8 cells from two independent assays is presented as the mean ± SEM (n=8; \*\*\**P*L<L0.001). (C), The left panel shows a magnified image of the IBs boxed in the merged panel shown in (A). The graphs (right panel) show the fluorescent intensity profiles along the indicated white lines drawn across one or more IBs. (D), IRF3 levels in HepG2 cells infected with EBOV trVLPs were analyzed by immunoblotting with an anti-IRF3 antibody at the indicated hours post infection (hpi). (E), The IRF3 intensity in cells infected with or without EBOV trVLPs for 48 h (the lower panel of (A)) was analyzed using ImageJ software. Differences between the two groups were evaluated using a two-sided unpaired Student’s *t*-test. The intensity of IRF3 in 5 cells from two independent assays is presented as the mean ± SEM (n = 5; ns, not significant).

### EBOV trVLPs infection attenuates the TBK1-IRF3 association and IRF3 nuclear translocation

Upon virus infection, IRF3, as a critical transcription factor in the IFN induction pathway, can be phosphorylated and activated by TBK1, and then phosphorylated IRF3 translocates from the cytoplasm into the nucleus, eliciting the expression of antiviral IFNs. Given the sequestration of IRF3 by EBOV trVLPs in IBs, the TBK1-IRF3 association in EBOV trVLPs-infected cells was assessed by an *in situ* Duolink proximity ligation assay (PLA). Cytoplasmic complexes consisting of endogenous TBK1 with IRF3 (the red signal) were observed in HepG2 cells treated with poly(I:C), which induces the activation of the RIG-I signal cascade and IRF3 phosphorylation, and poly(I:C)-induced TBK1: IRF3 complexes were significantly reduced by EBOV trVLPs infection (Fig. 3A and 3B). Decreased TBK1-IRF3 association was further demonstrated by immunoprecipitation (Fig. 3C). Moreover, as shown in Fig. 3D, Sendai virus (SeV) infection-induced IRF3 phosphorylation and nuclear translocation were significantly inhibited by EBOV trVLPs (Fig. 3D and Fig. 4A, 4B). Importantly, IRF3 was also recruited into IBs-like compartments in the cytoplasm in the cells infected with live EBOV (Fig. 4C). These data collectively suggested that the EBOV-mediated sequestration of IRF3 in IBs blocks IRF3 phosphorylation and nuclear translocation in the TBK1-IRF3 signaling cascade, which is critical for IFN induction.

**Fig. 3.**
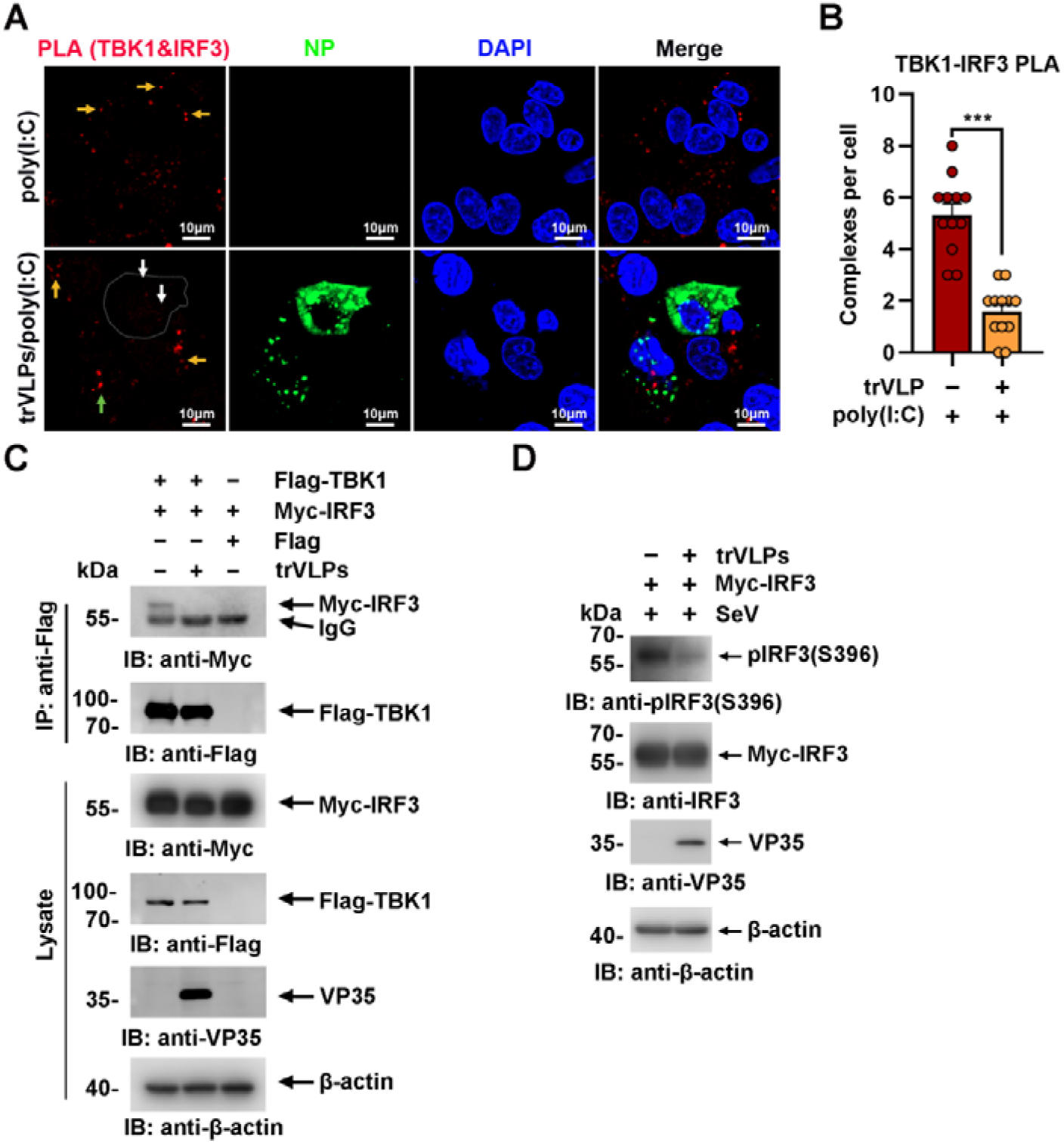
EBOV trVLPs inhibit IRF3 activation. (A), HepG2 cells were infected with or without the EBOV trVLPs. Thirty-six hours after infection, the cells were treated with or without 5 μg/ml poly(I:C) for 12 h and then subjected to *in situ* PLA with anti-TBK1 and anti-IRF3 antibodies and immunostaining with an anti-NP antibody (green). Nuclei were stained with DAPI (blue), and images were obtained using a Zeiss LSM 800 Meta confocal microscope. Arrows: white arrows indicate TBK1-IRF3 complexes in trVLPs-infected cells, and yellow and green arrows indicate TBK1-IRF3 complexes in uninfected and infected cells with small IBs, respectively. Scale bar, 10 μm. (B), The signal for the PLA complex in each cell in (A) was counted from at least 12 cells and is presented as the mean ± SEM. (C), Lysates of HEK293 cells cotransfected with or without the EBOV minigenome (p0) and the indicated plasmids were subjected to anti-Flag immunoprecipitation and analyzed by immunoblotting. (D), HEK293 cells were cotransfected with or without the EBOV minigenome (p0) and Myc-IRF3 plasmids. Thirty-six hours after transfection, the cells were infected with SeV at an MOI of 2 for 12 h, and the phosphorylation of IRF3 was analyzed by immunoblotting with an anti-IRF3-S396 antibody.

**Fig. 4.**
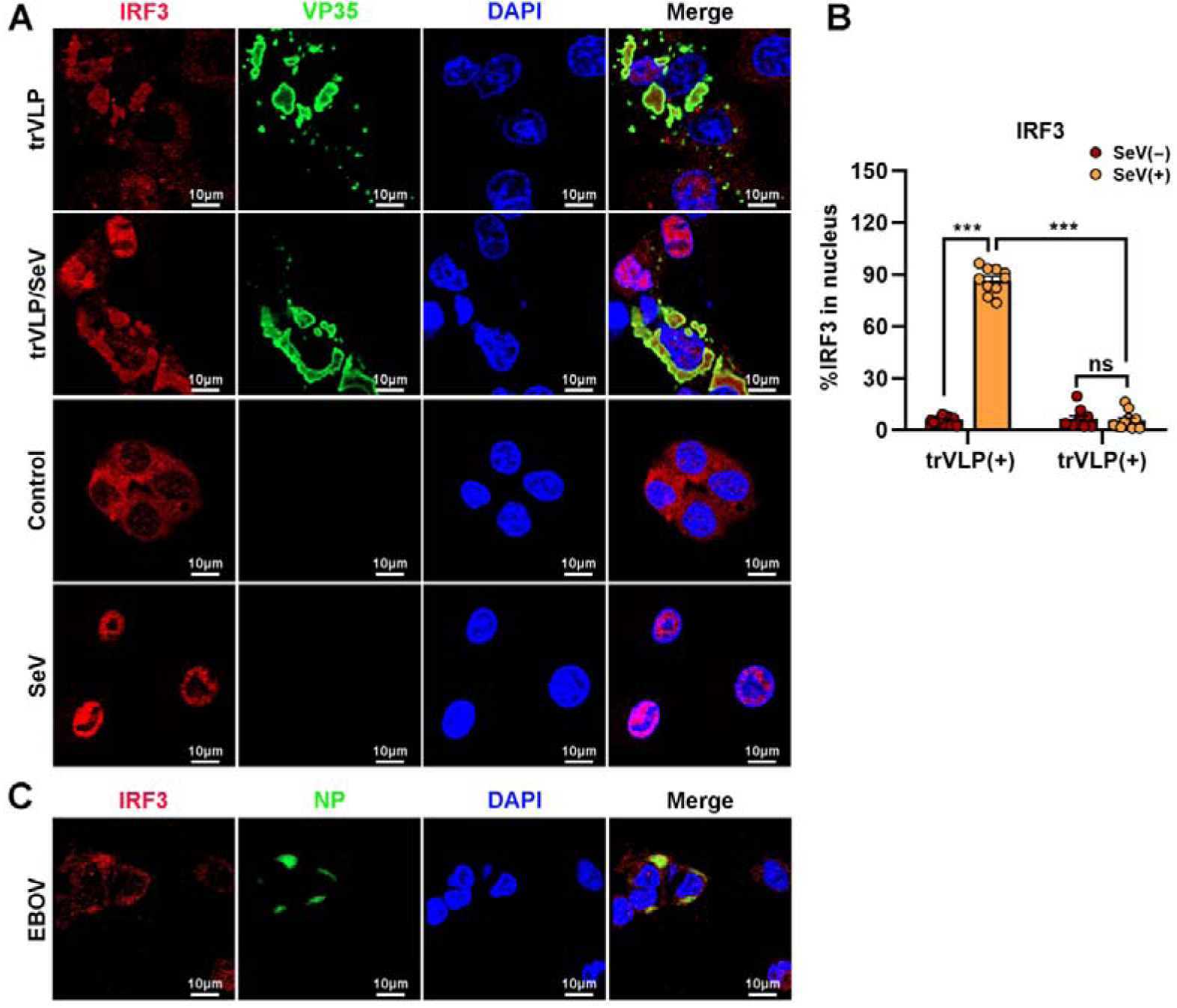
EBOV trVLPs inhibit nuclear translocation of IRF3. (A), HepG2 cells were infected with or without the EBOV trVLPs for 36 h, and the cells were infected with or without SeV at an MOI of 2 for another 12 h. The cells were then fixed and immunostained with anti-IRF3 (red) and anti-VP35 (green) antibodies. Nuclei were stained with DAPI (blue), and images were obtained using a Zeiss LSM 800 Meta confocal microscope. Scale bar, 10 μm. (B), The percentage of IRF3 nuclear distribution in (A) was analyzed using ImageJ software. The ratio of IRF3 distribution in ten cells from two independent assays is presented as the mean ± SEM (ns, not significant, \*\*\**P* < 0.001). (C), HepG2 cells infected with live EBOV (MOI=10) for 72 h were immunostained with anti-IRF3 (red) and anti-NP (green) antibodies. Nuclei were stained with DAPI (blue), and images were obtained using a Zeiss LSM 800 Meta confocal microscope. Scale bar, 10 μm.

### IBs-like structures formed by the viral proteins VP35 and NP play a key role in inducing IRF3 sequestration

Ectopic expression of NP alone (Noda, Watanabe, Sagara, & Kawaoka, 2007) or NP and the VP35 protein (Noda, Kolesnikova, Becker, & Kawaoka, 2011) in cells was sufficient to form IBs-like structures. To investigate the viral protein(s) involved in the sequestration of IRF3 in IBs, HepG2 cells were transiently transfected with plasmids encoding NP/VP35, NP/VP35/L, NP/VP35/L/VP30, NP/VP35/L/VP30/VP24, or NP/VP35/L/VP30/vRNA-RLuc/T7 and stained with anti-IRF3 and anti-NP at 48 hpi. Coexpression of NP and VP35 resulted in substantial sequestration of IRF3 in the IBs-like structure, which in turn resulted in a significant reduction of IRF3 in the nucleus, as observed in the cells transfected with vectors only and treated with poly(I:C) (Fig. 5A and 5B). Little if any VP35 or NP was demonstrated to interact with IRF3 by immunoprecipitation (Fig. 5 – figure supplement 1A-B). Compared to NP/VP35 coexpression, the presence of protein L, VP30 and VP24 showed little, if any, effects on IBs-like structure formation, IRF3 sequestration and nuclear IRF3 levels (Fig. 5A, 5B and Fig. 5 – figure supplement 2A-B). These results suggested that IBs-like structures as well as VP35 expression were indispensable for IRF3 sequestration.

**Fig. 5.**
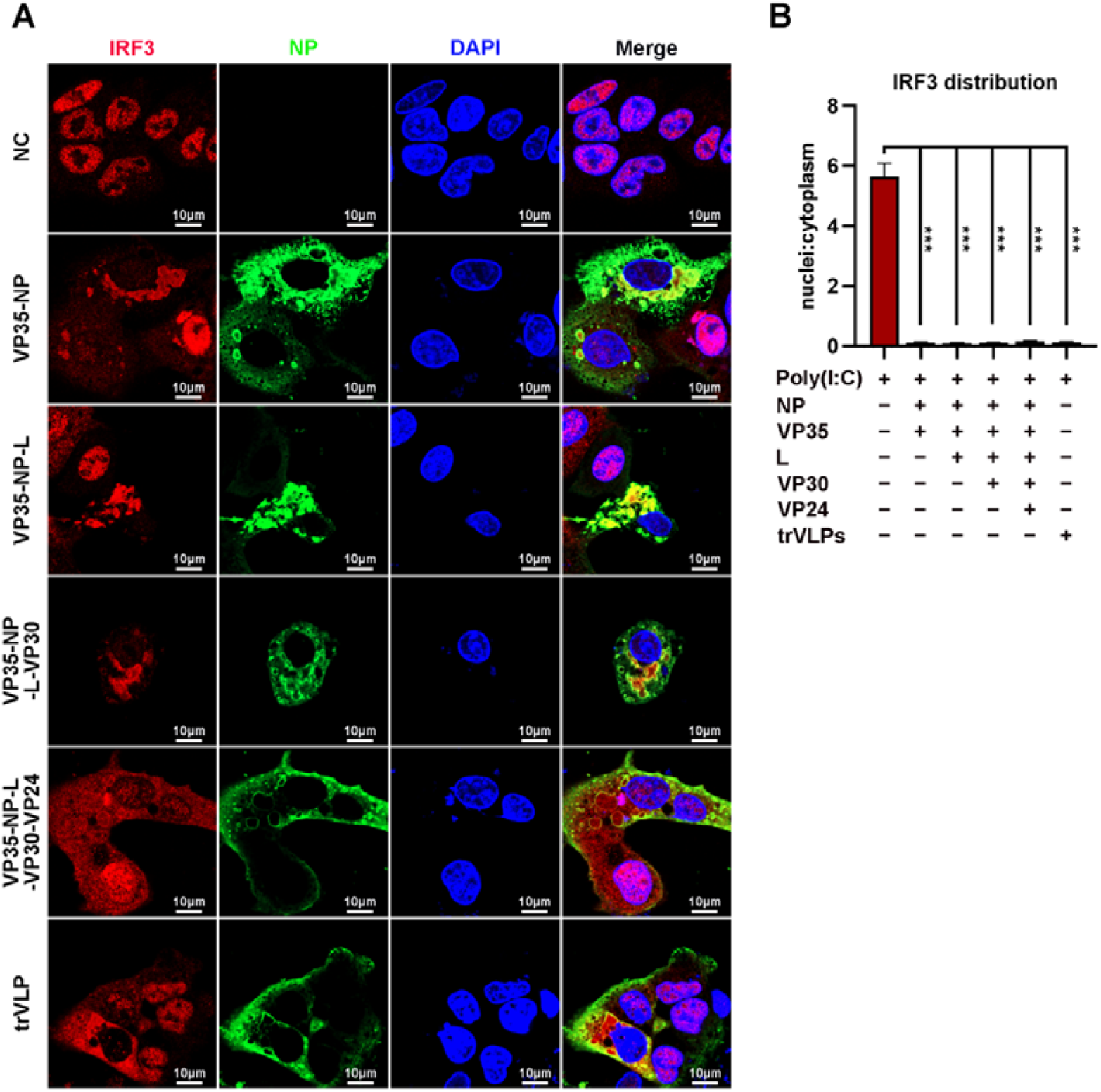
EBOV NP and VP35 play an important role in sequestering IRF3 into IBs. (A), HepG2 cells were transfected with the indicated plasmids for 36 h, and the cells were treated with 5 μg/ml poly(I:C) for another 12 h. Then, the cells were fixed and immunostained with anti-IRF3 (red) and anti-NP (green) antibodies. Nuclei were stained with DAPI (blue), and images were obtained using a Zeiss LSM 800 Meta confocal microscope. Scale bar, 10 μm. (B), The nuclear/cytoplasmic distribution of IRF3 in (A) was analyzed by ImageJ software. Differences between the two groups were evaluated using a two-sided unpaired Student’s *t*-test. The ratio of IRF3 distribution in at least 5 cells from two independent assays is presented as the mean ± SEM (n=5; \*\**P* < 0.01, \*\*\**P* < 0.001).

### VP35: STING interactions play an important role in isolating IRF3 into viral IBs

TBK1 and IKKε were spatially separated from VP35 upon infection by EBOV trVLPs (Fig. 1C and 1E), and IRF3 itself was demonstrated not to interact with VP35 and NP (Fig. 5 – figure supplement 1A-B), implying that other IRF3-interacting proteins might be involved in IRF3 sequestration in IBs upon viral infection. Stimulator of IFN genes (STING), an endoplasmic reticulum adaptor associated with IRF3 (Petrasek et al., 2013), was observed to interact with VP35 (Fig. 6A) and be recruited into IBs when the cells were infected by EBOV trVLPs (Fig. 6B and 6C). A substantial portion of STING was found to be recruited into IBs at 36 hpi in EBOV trVLPs-infected cells (Fig. 6D, 6E and Fig. 6 – figure supplement 1). STING knockdown by siRNA inhibited IRF3 sequestration in viral IBs (Fig. 6F and 6G). These results suggested that STING played important roles in the sequestration of IRF3 in viral IBs, possibly by interacting with VP35.

**Fig. 6.**
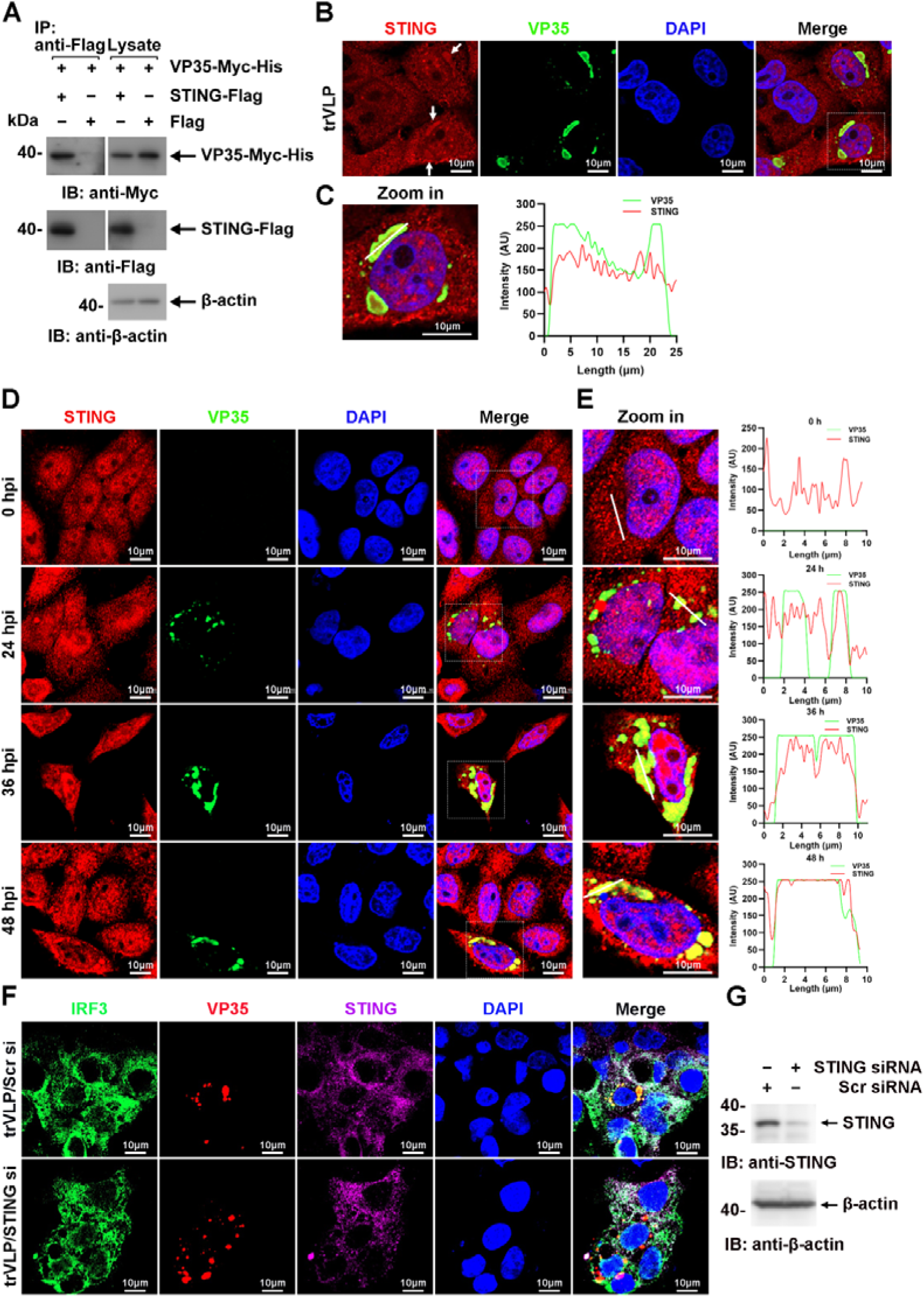
EBOV trVLPs recruit IRF3 into viral IBs via STING. (A), Lysates of HEK293 cells transfected with the indicated plasmids were subjected to anti-Flag immunoprecipitation and analyzed by immunoblotting. (B) HepG2 cells were transfected with the EBOV minigenome (p0). Forty-eight hours after infection, the cells were fixed and immunostained with anti-STING (red) and anti-VP35 (green) antibodies. White arrows: STING in IBs. Nuclei were stained with DAPI (blue), and images were obtained using a Zeiss LSM 800 Meta confocal microscope. Scale bar, 10 μm. (C) The left panel shows a magnified image of the IBs boxed in the merged panel of (B). The graphs (right panel) show the fluorescent intensity profiles along the indicated white lines drawn across one or more IBs. (D) HepG2 cells were infected with the EBOV trVLPs. At the indicated hours post infection (hpi), cells were fixed and immunostained with anti-STING (red) and anti-VP35 (green) antibodies. Nuclei were stained with DAPI (blue), and images were obtained using a Zeiss LSM 800 Meta confocal microscope. Scale bar, 10 μm. The data from two independent replicates are presented. (E) The left panel shows a magnified image of the IBs boxed in the merged panel of (D). The graphs (right panel) show fluorescent intensity profiles along the indicated white lines drawn across one or more IBs. (F and G) HepG2 cells were transfected with STING siRNA (STING si) or scrambled siRNA (Scr si) for 6 h. The cells were then infected with the EBOV trVLPs for 36 h and then immunostained with Fluor 488-conjugated-anti-IRF3 (green), anti-VP35 (red) and anti-STING (purple) antibodies. Nuclei were stained with DAPI (blue), and images were obtained using a Zeiss LSM 800 Meta confocal microscope. Scale bar, 10 μm. The silencing efficiency of STING siRNA was determined by immunoblotting (G).

### Viral IBs-induced IRF3 sequestration suppresses IFN-**β** production

EBOV trVLPs could hijack IRF3 and sequester IRF3 into IBs and thus block the nuclear translocation of IRF3, which suggested that EBOV trVLPs may suppress IRF3-driven IFN-β production. As reported previously (Basler et al., 2000), expression of VP35 (Fig. 7A), but not NP, resulted in a mild inhibition of SeV-induced IFN-β-Luc expression (Fig. 7B). Coexpression of VP35 and NP, which led to the formation of IBs and the sequestration of IRF3 (Fig. 5A), suppressed IFN-β-Luc expression much more potently than VP35 expression alone (Fig. 7B). Coexpression of NP/VP35/L/VP30 was more potent in the inhibition of SeV-induced IFN-β-Luc expression than NP/VP35 (Fig. 7B). Moreover, coexpression of NP/VP35/VP30/L almost completely suppressed poly(I:C)-induced IFN-β transcription (Fig. 7C). *IRF3* depletion showed little, if any, effect on IFN-β transcription upon NP/VP35/L/VP30 coexpression (Fig. 7C and Fig. 7 – figure supplement 1A), which suggested that NP/VP35/L/VP30 coexpression was similarly powerful as *IRF3* depletion in antagonizing IFN-β expression. In wild-type cells but not *IRF3*-depleted cells, the coexpression of NP/VP35/L/VP30 had a significantly greater ability to inhibit SeV-induced transcription of IFN-β downstream genes, such as CXCL10, ISG15 and ISG56, than VP35 expression alone (Fig. 7D-F). These results strongly suggested that the sequestration of IRF3 in viral IBs was substantially more powerful than that upon VP35 expression.

**Fig. 7.**
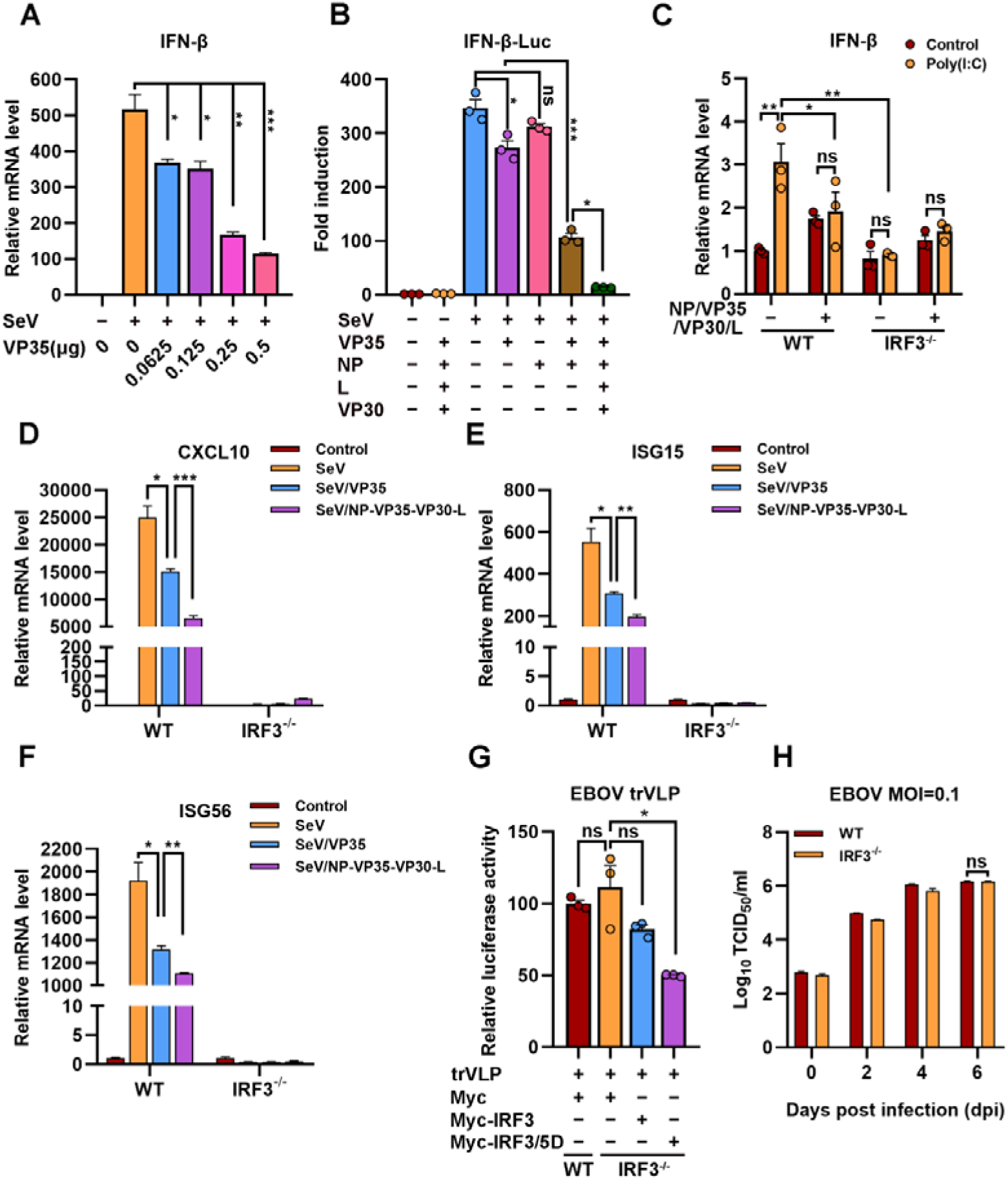
The hijacking of IRF3 by viral IBs inhibits IFN-β production. (A), HEK293 cells were transfected with the indicated plasmids for 24 h, and the cells were infected with or without SeV at an MOI of 2 for another 12 h. The mRNA level of IFN-β was quantified by qRT-PCR. Differences between the two groups were evaluated by a two-sided unpaired Student’s *t*-test. The data are presented as the means☐±☐SEM (\**P*☐<☐0.05, \*\**P*☐<☐0.01, \*\*\**P*☐<☐0.001). (B), HEK293 cells were cotransfected with the firefly luciferase reporter plasmid pGL3-IFN-β-Luc, the *Renilla* luciferase control plasmid pRL-TK, and viral protein expression plasmids (0.0625 μg of pCAGGS-NP, 0.0625 μg of pCAGGS-VP35, 0.0375 μg of pCAGGS-VP30 and 0.5 μg of pCAGGS-L) for 24 h, and the cells were infected with or without SeV at an MOI of 2 for another 12 h. The luciferase activities were then analyzed. The data were analyzed to determine the fold induction by normalizing the firefly luciferase activity to the *Renilla* luciferase activity. Empty plasmid without SeV infection was used as a control, and the corresponding data point was set to 100%. Differences between the two groups were evaluated using a two-sided unpaired Student’s *t*-test. The data are presented as the means☐±☐SEM (ns, not significant, \**P*☐<☐0.05, \*\*\**P*☐<☐0.001). (C), Wild-type (WT) and *IRF3*-depleted (IRF3^-/-^) HeLa cells were transfected with or without pCASSG-NP, pCASSG-VP35, pCASSG-VP30 and pCASSG-L plasmids for 36 h and then treated with or without 5 μg/ml poly(I:C) for 12 h. The mRNA level of IFN-β was quantified by qRT-PCR. Differences between the two groups were evaluated using a two-sided unpaired Student’s *t*-test. The data are presented as the means☐±☐SEM (ns, not significant, \**P*☐<☐0.05). (D-F), Wild-type (WT) and *IRF3*-depleted (IRF3^-/-^) HeLa cells were transfected with or without pCAGGS-VP35 or pCASSG-NP, pCASSG-VP35, pCASSG-VP30 and pCASSG-L plasmids for 36 h, and the cells were infected with or without SeV at an MOI of 5 for another 12 h. The mRNA level of CXCL10 (D), ISG15 (E) and ISG56 (F) were quantified by qRT-PCR. Differences between the two groups were evaluated using a two-sided unpaired Student’s *t*-test. The data are presented as the means☐±☐SEM (\**P*☐<☐0.05, \*\**P*☐<☐0.01, \*\*\**P*☐<☐0.001). (G), Wild-type (WT) and *IRF3* knockout (IRF3^-/-^) HeLa cells were transfected with the EBOV minigenome (p0), pGL3-promoter and Myc-vector, Myc-IRF3 or Myc-IRF3/5D plasmids for 96 h. The amounts of trVLPs were determined by a luciferase activity assay (left panel). Differences between the two groups were evaluated by a two-sided unpaired Student’s *t*-test. The data are presented as the means☐±☐SEM (ns, not significant, \*\*\**P*☐<☐0.001). (H). Wild-type (WT) and *IRF3*-knockout (IRF3^-/-^) HeLa cells were infected with live EBOV (MOI = 0.1). The cell culture supernatants were collected on the indicated days post infection (dpi), and the viral titers were quantified as TCID_50_ by a plaque assay. Differences between the two groups were evaluated using a two-sided unpaired Student’s *t*-test. The data are presented as the means☐±☐SEM (ns, not significant).

We next assessed the effect of IRF3 hijacking and sequestration by viral IBs on EBOV trVLPs replication. Compared with wild-type cells, *IRF3* depletion showed little, if any, effect on EBOV replication, as indicated by luciferase activity, suggesting that trVLPs efficiently blocked IRF3 signaling (Fig. 7G and Fig. 7 – figure supplement 1B). Moreover, the overexpression of IRF3/5D (a phospho-mimic of activated IRF3), but not IRF3, inhibited EBOV trVLPs replication in *IRF3*-depleted cells (Fig. 7G). Importantly, compared with wild-type cells, *IRF3* depletion showed little, if any, effect on EBOV replication in the cells infected with live EBOV (Fig. 7H). Taken together, these results suggest that the hijacking of IRF3 and sequestration into IBs by EBOV can be significantly more potent in the inhibition of IFN-I production and thereby antagonizes the inhibitory effect of IFN-I on viral replication.

## Discussion

Accumulating evidence suggests that EBOV has established multiple ways to antagonize host innate immune responses to maintain viral replication. Several EBOV proteins (VP35, VP24, GP, VP30 and VP40) are known to participate in host immune evasion to facilitate viral replication and pathogenesis (Audet & Kobinger, 2015; Bhattacharyya, 2021; Cantoni & Rossman, 2018). VP35 was demonstrated to suppress IFN-I production by inhibiting IRF3/7 phosphorylation, disrupting DC maturation, and facilitating the escape of immune sensation by dsRNA (Basler, 2015; Cardenas et al., 2006; Messaoudi, Amarasinghe, & Basler, 2015; Prins et al., 2009). VP30 and VP40 suppress RNA silencing by interacting with Dicer and modulating RNA interference components via exosomes, respectively (Fabozzi, Nabel, Dolan, & Sullivan, 2011; Pleet, DeMarino, Lepene, Aman, & Kashanchi, 2017). VP24 and GP are also known to block IFN-I signaling by hiding MHC-1 on the cell surface and counteracting tetherin or interfering with established immune responses by adsorbing antibodies against GP, respectively (Audet & Kobinger, 2015; Bhattacharyya, 2021).

Viral IBs are a characteristic of cellular EBOV infection and are important sites for viral RNA replication, and NP and VP35 are extremely critical proteins for the formation of IBs (Hoenen et al., 2012). However, whether viral IBs are involved in antagonizing IFN-I production during EBOV trVLPs infection has not yet been reported. Here, we found that IRF3 is hijacked and sequestered into EBOV IBs by viral infection (Fig. 1A), which demonstrates that viral IBs are utilized for IRF3 compartmentalization. Meanwhile, this compartmentalization resulted in the spatial isolation of IRF3 from the kinases TBK1 and IKKε (Fig. 1C and 1E). This suggests that IRF3 deprivation by viral IBs may antagonize host antiviral signaling by inhibiting IFN-I production signaling.

As expected, the expression of NP/VP35/VP30/L, which is involved in the composition of IBs, was significantly more antagonistic to SeV-induced IFN-β production than the expression of VP35 alone (Fig. 7B). In addition, the expression of NP/VP35/VP30/L can significantly antagonize the promoting effect of poly(I:C) on IFN-β transcription, and IRF3 knockout could not further inhibit the transcription of IFN-β (Fig. 7C), which may be because viral hijacking of IRF3 into IBs nearly completely antagonized its function of promoting IFN-β production. In this study, the effect of poly(I:C) is consistent with the results obtained with SeV, which indicates that poly(I:C) may mainly activate the RLR signaling pathway (Fig. 3A). As shown in Fig. 7D-F, the expression of NP/VP35/VP30/L significantly inhibited the ability of SeV to promote the transcription of IFN-β downstream genes (CXCL10, ISG15 and ISG56) but did not completely suppress the effect of SeV, which may be due to the low transfection rate of HeLa cells. Furthermore, the knockout of IRF3 in cells could not further promote EBOV and EBOV trVLPs replication compared with that observed in wild-type cells (Fig. 7G and 7H), which may have been because IRF3 was hijacked into viral IBs and could not be phosphorylated into the nucleus to regulate IFN-I production. These results suggest that viral IBs act as virus-built ‘jails’ to imprison transcription factors and present a novel and possible common mechanism of viral immune evasion in which the critical signaling molecule IRF3 is spatially segregated from the antiviral kinases TBK1 and IKKε.

Although almost all IRF3 could be sequestered to viral IBs formed by VP35 and NP (Fig. 5A and 5B), we found that neither VP35 nor NP interacted with IRF3 (Fig. 5 – figure supplement 1). Here, we found that VP35 interacts with STING and colocalizes in IBs and that knockdown of STING inhibits the sequestration of IRF3 in IBs (Fig. 6A-G). These results suggest that VP35 may hijack IRF3 into IBs through STING. However, whether other host proteins are involved in this process and the role of NP in the recruitment of IRF3 by VP35 remain unclear. In addition, we found that VP35 may hijack IRF3 into IBs via STING association (Fig. 6A-C); however, whether VP35 activates the STING-IRF3 pathway in a cGAS-independent manner by interacting with STING and the molecular mechanism remain to be further investigated.

In summary, EBOV VP35 sequesters IRF3 into viral IBs and inhibits the association of IRF3 with TBK1 and IKKε, preventing IRF3 from entering the nucleus and thereby inhibiting IFN-I production (Fig. 8). Therefore, this study reveals a new strategy by which EBOV escapes the innate immune response and provides new ideas for Ebola virus disease treatment.

**Fig. 8.**
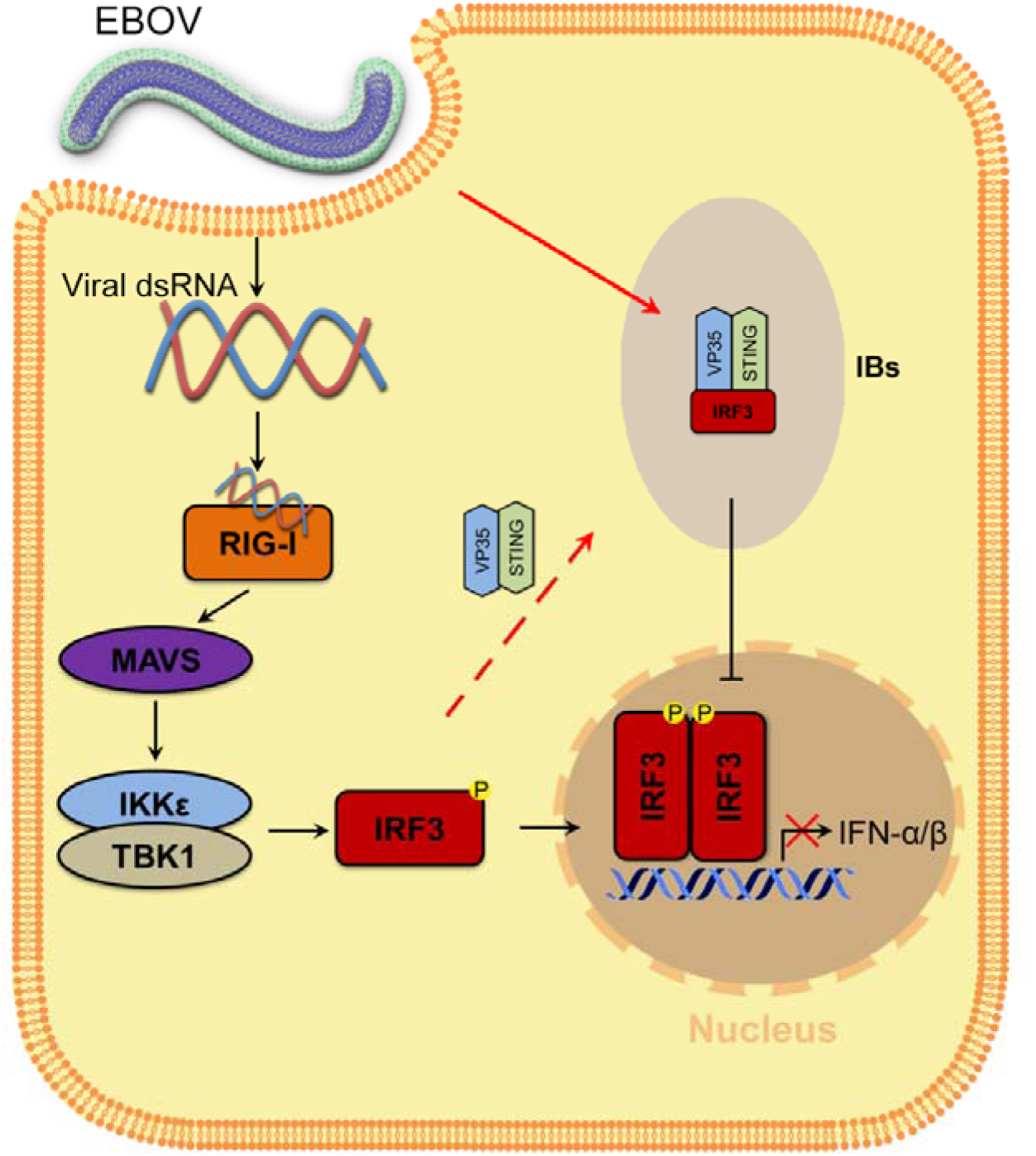
Model of the molecular mechanism by which EBOV hijacks IRF3 into viral IBs through VP35-STING to comprehensively disrupt IFN-I production. VP35 sequesters IRF3 to EBOV IBs, which in turn spatially segregates IRF3 from TBK1 and IKKε, blocks RLR signaling and inhibits IFN-I production.

## Materials and Methods

### Cell lines and transfections

HEK293, HeLa and IRF3-knockout HeLa cells (ABclonal, RM02113) were grown in Dulbecco’s modified Eagle’s medium (DMEM, Gibco). HepG2 cells were grown in minimum essential medium (MEM, Gibco) supplemented with a 1% nonessential amino acid solution (NEAA, Gibco). All media were supplemented with 10% heat-inactivated fetal bovine serum (Gibco), 2 mM L-glutamine, 100 units/ml penicillin and 100 units/ml streptomycin, and cells were grown at 37°C under an atmosphere with 5% CO_2_. Transient transfection was performed with Lipofectamine 3000 (Invitrogen) according to the manufacturer’s instructions.

### Vectors and viruses

Flag-tagged VP35, NP, STING, TBK1 and IRF3 vectors were constructed by cloning the corresponding gene fragments into a pcDNA3.0-based Flag-vector (Invitrogen). Myc-tagged VP35, IRF3 and IRF3/5D vectors were constructed by inserting the corresponding gene fragments into the pCMV-Myc vector (Clontech). All the constructs were validated by Sanger DNA sequencing.

SeV was amplified in 9- to 11-day embryonated specific pathogen-free (SPF) eggs. Live EBOV (Mayinga strain) is preserved by the BSL-4 Lab at the Wuhan Institute of Virology, Chinese Academy of Sciences.

### Immunoprecipitation and immunoblot analysis

Cell lysates were prepared in lysis buffer containing 1% Nonidet P-40 and protease inhibitor cocktail (Roche) (Cao, Leng, & Kufe, 2003). Soluble proteins were immunoprecipitated using anti-Flag (M2, Sigma), anti-Myc (Sigma) or IgG of the same isotype from the same species as a negative control (Sigma). An aliquot of the total lysate (5%, v/v) was included as a control. Immunoblotting was performed with horseradish peroxidase (HRP)-conjugated anti-Myc (Sigma), HRP-conjugated anti-Flag (Sigma), HRP-conjugated anti-β-actin (Sigma), anti-VP35 (Creative Diagnostics), anti-IRF3 (Cell Signaling Technology), anti-STING (Proteintech) or anti-NP (Sino Biological) antibodies. The antigen-antibody complexes were visualized via chemiluminescence (Immobilon Western Chemiluminescent HRP Substrate, Millipore). A PageRuler Western marker (Thermo) was used as a molecular weight standard.

### Gene silencing using siRNA

For gene knockdown in HepG2 cells, cells maintained in 6-well plates were transfected with 100 pmol STING small interfering RNA (siRNA) (sense, 5’- GCACCUGUGUCCUGGAGUATT -3’; antisense, 5’-UACUCCAGGACACAGGUGCTT -3’) or the same concentration of scrambled siRNA (sense, 5’- UUCUCCGAACGUGUCACGUTT -3’; antisense, 5’- ACGUGACACGUUCGGAGAATT -3’) purchased from Tsingke Biotechnology (Beijing, China) with Lipofectamine 3000 (Invitrogen) according to the manufacturer’s recommendations.

### Reverse transcription and quantitative RT-PCR

Total cellular RNA was prepared using an RNeasy Mini kit (QIAgen, USA) according to the manufacturer’s protocol. For cDNA synthesis, 0.5 μg of RNA was first digested with gDNA Eraser to remove contaminated DNA and then reverse transcribed using ReverTra Ace qPCR RT Master Mix with gDNA Remover (FSQ-301, Toyobo) in a 20 μL reaction volume. Then, 1 μL of cDNA was used as a template for quantitative PCR. The following primers were used in these experiments:

h-IFN-β F: 5’-AGGACAGGATGAACTTTGAC-3’

h-IFN-β R: 5’-TGATAGACATTAGCCAGGAG-3’;

h-CXCL10-F: 5’-TCCCATCACTTCCCTACATG-3’;

h-CXCL10-R: 5’-TGAAGCAGGGTCAGAACATC-3’;

h-ISG15-F: 5’-TCCTGGTGAGGAATAACAAGGG-3’;

h-ISG15-R: 5’-CTCAGCCAGAACAGGTCGTC-3’;

h-ISG56-F: 5’-TCGGAGAAAGGCATTAGATC-3’;

h-ISG56-R: 5’-GACCTTGTCTCACAGAGTTC-3’;

h-GAPDH F: 5’-AAggTCATCCCTgAgCTgAAC-3’;

h-GAPDH R: 5’-ACgCCTgCTTCACCACCTTCT-3’.

The samples were denatured at 95°C for 2 min, followed by 40 cycles of amplification (15 s at 94°C for denaturation, 60 s at 60°C for annealing and extension). Quantitative RT-PCR (qRT-PCR) was performed using SYBR Green Real-time PCR Master Mix (QPK-201, Toyobo) with the QuantStudio 6 Flex multicolor real-time PCR detection system (ABI). Relative mRNA levels were normalized to GAPDH levels and calculated using the 2^-ΔΔ*CT*^ method (Livak & Schmittgen, 2001). The means (upper limit of the box) ± SEM (error bars) of three independent experiments are presented in the figures.

### *In situ* proximity ligation assay

Duolink *in situ* PLA (Sigma) was used to detect the endogenous association of IRF3 and TBK1 in cells. In brief, HepG2 cells plated on glass coverslips were transfected with EBOV minigenome plasmids. After fixation with 4% formaldehyde, the cells were permeabilized with 0.3% Triton X-100 in PBS for 15 min. After blocking with blocking buffer (Sigma, DUO82007), the cells were incubated with mouse anti-IRF3 (Cell Signaling Technology) and rabbit anti-TBK1 (Abcam) primary antibodies. The nuclei were stained with DAPI (blue). The red fluorescent spots generated from the DNA amplification-based reporter system combined with oligonucleotide-labeled secondary antibodies were detected with a Zeiss LSM 800 Meta confocal microscope (Carl Zeiss).

### Immunofluorescence microscopy

Cells were transfected, fixed, permeabilized and blocked as described above. Then, after incubation with anti-TBK1 (Cell Signaling Technology), anti-IKKε (Cell Signaling Technology), anti-IRF3 (Cell Signaling Technology), anti-VP35 (Creative Diagnostics), anti-NP (Sino Biological), or anti-STING (Bioss) antibodies overnight at 4°C, the cells were washed three times with PBST buffer and then incubated with 488-conjugated anti-IRF3 (Proteintech) antibodies, FITC- or TRITC-conjugated goat anti-rabbit (or anti-mouse) IgG secondary antibodies for another 1 h at room temperature. The cells were then stained with DAPI after washing and imaged using a laser scanning confocal microscope (Zeiss LSM 800 Meta) with a 63× oil immersion lens.

### Luciferase reporter assay

The IFN-I production assay was performed as described previously (Zhu et al., 2022). Briefly, HEK293 cells (1×10^5^ cells per well in a 24-well plate) were cotransfected with the indicated amount of pCAGGS-NP (62.5 ng)/pCAGGS-VP35 (62.5 ng)/pCAGGS-VP30 (37.5 ng)/pCAGGS-L (500 ng), 200 ng of the IFN-β reporter plasmid (Promega, USA) and 4 ng of *Renilla* luciferase plasmid. An empty vector was used to ensure that each well contained the same plasmid concentration. After 24 h, the cells were treated with SeV (MOI=2) or 5 μg/ml poly(I:C) for 12 h, and the luciferase activity of the cell lysates was analyzed with the dual-luciferase reporter assay system (Promega, E1960) using a GloMax 20/20 luminometer (Promega, USA). Values were obtained by normalizing the luciferase values to the Renilla values. Fold induction was determined by setting the results from the group transfected with vector without Flag-VP35 to a value of 1.

### EBOV trVLPs assay

The replication of EBOV in the cells was evaluated with the minigenome system (Hoenen et al., 2014). Briefly, producer cells (p0) were cotransfected with p4cis-vRNA-RLuc (250 ng) and pCAGGS-T7 (250 ng) for T7 RNA polymerase expression and 4 plasmids for EBOV protein expression (pCAGGS-NP (125 ng), pCAGGS-VP35 (125 ng), pCAGGS-VP30 (75 ng), and pCAGGS-L (1,000 ng)), as well as the luciferase reporter vector pGL3-Promoter (Youbio, 25 ng). One day after transfection, the medium was replaced with medium containing 5% FBS, and the cells were then incubated for another 3 days. Viral replication was determined by intracellular luciferase activities using a dual-luciferase reporter assay kit (Promega, E1960) after cell lysis with passive lysis buffer (PLB, Promega). For immunofluorescence experiments, cells were harvested 48 h after transfection.

### Transmission electron microscopy (TEM)

HepG2 cells transfected with EBOV minigenome p0-related plasmids were washed with PBS, fixed with 2.5% glutaraldehyde, and then prestained with osmium tetroxide. Eighty-nanometer-thick serial sections were then cut and stained with uranyl acetate and lead citrate. Images were acquired with a transmission electron microscope (Hitachi, H-7650) operating at 80 kV.

### EBOV infection assay

HepG2, HeLa or *IRF3*-depleted HeLa cells grown to ∼70% confluency in 12-well plates (for viral proliferation) or 12-well plates with a 18-mm coverslip (for immunofluorescence microscopy) were incubated with the EBOV Mayinga strain, which was tittered in Vero E6 cells, at 37°C for 1 h at the indicated MOI. Then, the cells were washed 3 times with PBS, and fresh medium was added to the cells, which were incubated at 37°C for 72 h (for microscopy) or the indicated times (0 days, 2 days, 4 days and 6 days; for the viral proliferation assay). Subsequently, the cells on the coverslip were fixed with 4% formaldehyde for immunofluorescent straining, and the supernatants were collected at the indicated times for viral titration following the requirements of the BSL-4 laboratory. The viral titers were determined by plaque formation assay. Briefly, 10-fold serially diluted samples (100 μl) were added to 96-well plates containing 1×10^4^ Vero E6 cells per well and incubated for 1 h at 37°C in a 5% CO_2_ incubator. Then, 100 μl of medium containing 2% FBS was added to each well. After incubation for 5–7 days at 37°C in a 5% CO_2_ incubator, the cytopathic effect (CPE) was observed, and the median tissue culture infective dose (TCID_50_)/ml was calculated. All work with live EBOV was performed with BSL-4 containment.

### Statistical analyses

Graphical representation and statistical analyses were performed using Prism 8 software (GraphPad Software). Unless indicated otherwise, the results are presented as the means (upper limit of the box) ± SEM (error bars) from three independent experiments conducted in duplicate. An unpaired two-tailed *t*-test was used for the analysis of two groups. Data were considered significant when *P* < 0.05 (*), *P* < 0.01 (**), and *P* < 0.001 (***).

### Data availability

All the data shown in this paper are available from the corresponding authors upon reasonable request.

## Acknowledgments

This work was supported by the National Natural Science Foundation of China (82372255), the Advanced Customer Cultivation Project of the Wuhan National Biosafety Laboratory, the Chinese Academy of Sciences [2022ACCP-MS04] and the National Major Science and Technology Projects of China [2018ZX09711003-005-005 to T.G. and 2022YFC2600704 to H.L.].

## Author Contributions

L.Z., X.L., and C.C. designed and supervised the study. L.Z., J.J., and T.W. performed the experiments. Y.H. performed the experiment related to EBOV infection in the BSL-4 laboratory. T.G., H.L., Q.D., Y.Hu, Y.J., Z.L, and P.L. analyzed the data. L.Z, X.L., and C.C. wrote the manuscript. All authors have read and approved the article.

---

**Fig. 1 – figure supplement 1.**
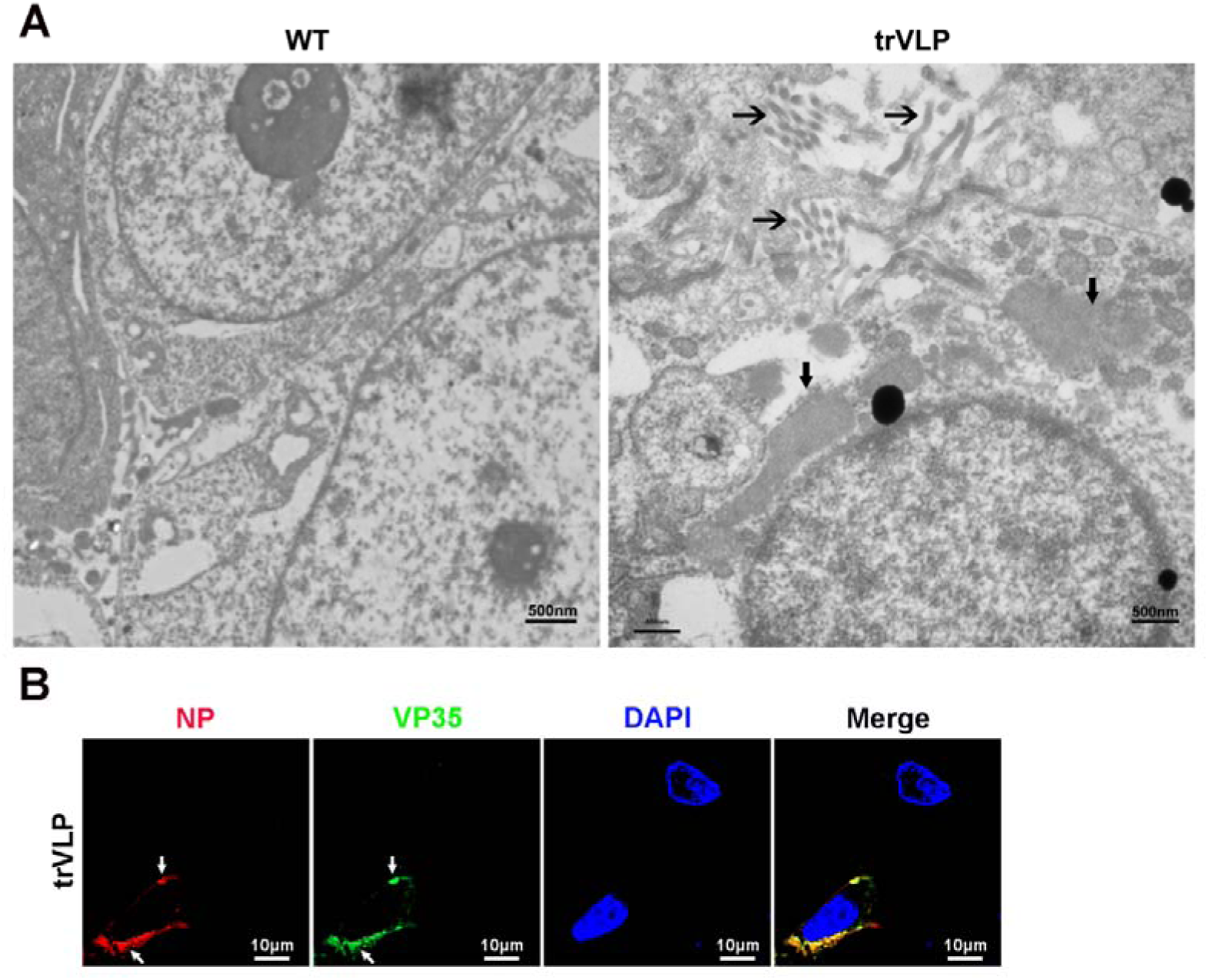
Transmission electron microscopy and immunofluorescence detection of EBOV trVLPs and IBs. (A), HepG2 cells infected with or without EBOV trVLPs were fixed and observed with a HITACHI H-7650 transmission electron microscope at an accelerating voltage of 80 kV. The IBs and viral particles (right panel) are marked with bold arrows and regular arrows, respectively. Scale bar, 500 nm. (B), HepG2 cells infected with the EBOV trVLPs were immunostained with anti-NP (red) and anti-VP35 (green) antibodies. Nuclei were stained with DAPI (blue), and images were obtained using a Zeiss LSM 800 Meta confocal microscope. Scale bar, 10 μm.

**Fig. 5 – figure supplement 1.**
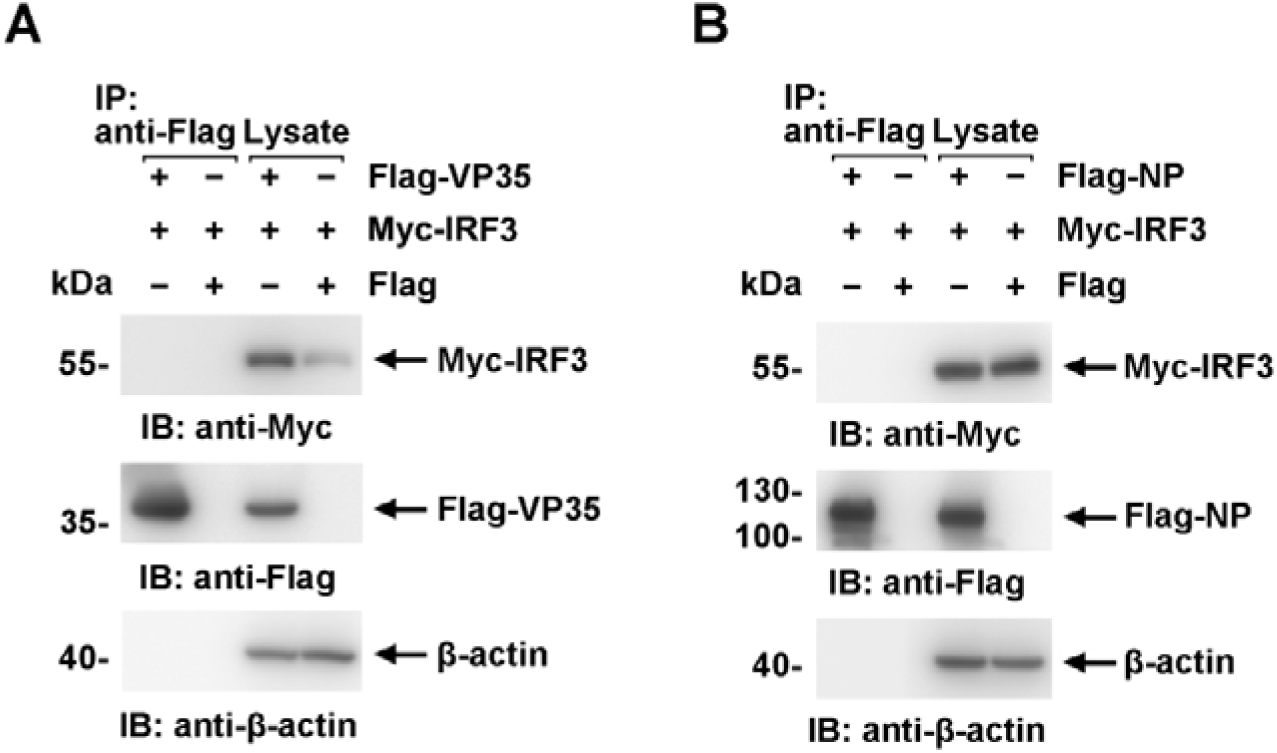
Neither VP35 nor NP interacts directly with IRF3 in cells. (A and B) Lysates of HEK293 cells transfected with the indicated plasmids were subjected to anti-Flag immunoprecipitation and analyzed by immunoblotting. The data from two independent replicates are presented.

**Fig. 5 – figure supplement 2.**
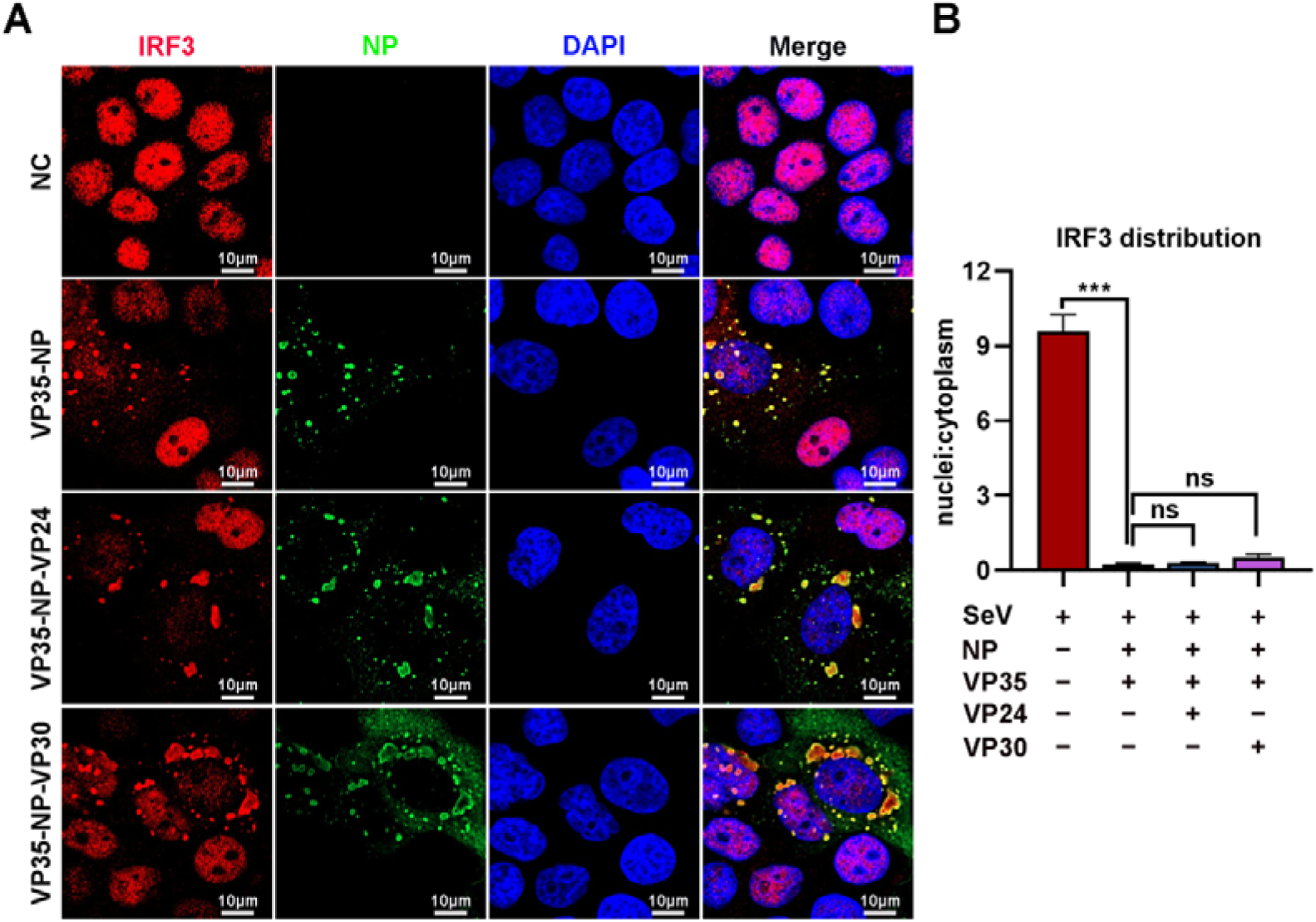
Neither VP24 nor VP30 plays an important role in sequestering IRF3 into IBs. (A), HepG2 cells were transfected with the indicated plasmids for 36 h, and the cells were infected with SeV at an MOI of 2 for another 12 h. The cells were then fixed and immunostained with anti-IRF3 (red) and anti-NP (green) antibodies. Nuclei were stained with DAPI (blue), and images were obtained using a Zeiss LSM 800 Meta confocal microscope. Scale bar, 10 μm. (B), The nuclear/cytoplasmic distribution of IRF3 in (A) was analyzed using ImageJ software. Differences between the two groups were evaluated using a two-sided unpaired Student’s *t*-test. The ratio of IRF3 distribution in at least 5 cells from two independent assays is presented as the mean ± SEM (n = 5; ns, not significant, \*\*\**P* < 0.001).

**Fig. 6 – figure supplement 1.**
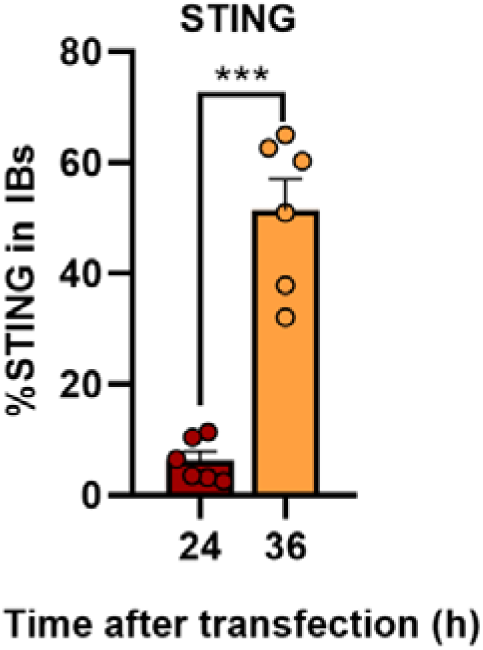
EBOV trVLPs recruit STING into viral IBs. The percentage of STING distribution in IBs at different time points in cells infected with EBOV trVLPs in Fig. 6D was analyzed with R programming language. The intensity of STING in 6 cells from two independent assays is presented as the mean ± SEM (n=6; \*\*\**P*☐<☐0.001).

**Fig. 7 – figure supplement 1.**
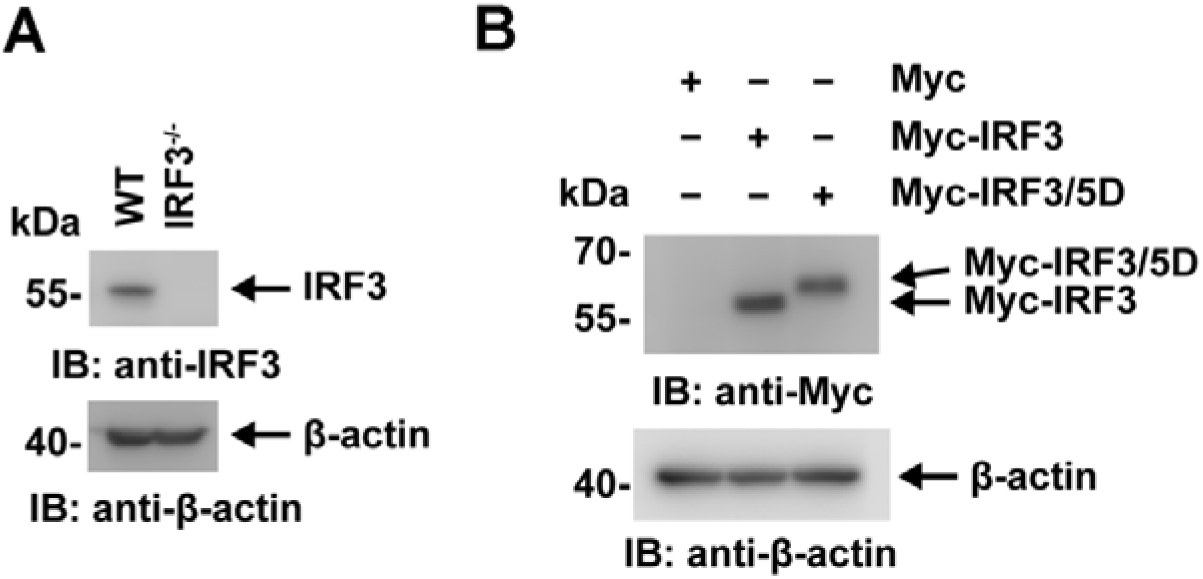
The expression of IRF3 and its mutants were detected by immunoblotting. (A) Lysates of WT and IRF3^-/-^ HeLa cells were analyzed by immunoblotting with an anti-IRF3 antibody. (B) Lysates of WT and IRF3^-/-^ HeLa cells transfected with Myc-vector, Myc-IRF3 or Myc-IRF3/5D were analyzed by immunoblotting with the indicated antibodies.

